# Multiscale modeling predicts dependence of mesenchymally transitioned tumor niche fitness on cell-cell and cell-matrix adhesions

**DOI:** 10.1101/2025.06.25.661486

**Authors:** C. Venkata Sai Prasanna, Mohit Kumar Jolly, Ramray Bhat

**Affiliations:** IISc Mathematics Initiative PhD Program, Indian Institute of Science, Bangalore, 560012, India; Department of Bioengineering, Indian Institute of Science, Bangalore, 560012, India; Department of Developmental Biology and Genetics, Indian Institute of Science, Bangalore, 560012, India

**Keywords:** **<u>Keywords</u>**: Epithelial-Mesenchymal Transition (EMT), tumor niche, cell-ECM adhesion, niche fitness, collective cancer invasion (CCI), niche domination

## Abstract

Invasion of cancer cells is often characterized by a transition in phenotype of cells or their niches from an epithelial to a mesenchymal state (EMT). Under what conditions do transitioned niches acquire greater fitness than, and outcompete, their parental un-transitioned niches, is not well-understood. Here, we use a Cellular Potts model-based multiscale computational framework to investigate this question. Inducing an EMT in a single cell at the edge of an early-growing tumor surrounded by a fibrillar extracellular matrix (ECM) allows us to temporally trace inter-niche competitions. We observe that the transitioned niche dominates the population it arises from and invades better when surrounded by dense ECM. An increase in cell-ECM adhesion by itself drives domination at 50% probability, such that the transitioned population invades faster and contributes further to collective invasion of the whole tumor. Decrease in inter- and intra-niche cell-cell adhesion by itself is not sufficient to achieve domination. However, added to increased cell-ECM adhesion, loss of intra-niche (but not inter-niche adhesion) restores the probability, but not the extent, with which domination by the transitioned niche is achieved by attenuating its confinement by its parental population. Our simulations reveal the forces regulating such confinement and how cell-cell and cell-ECM adhesion, stochastic invasion dynamics, and ECM density contribute nuancedly to distinct aspects of inter-niche competitions within tumor populations and their fitness.

## Introduction

Cancer metastasis is a complex process involving detachment of cancer cells from primary tumor as single cells or clusters, entry into circulation or intravasation, survival in the bloodstream, and colonization of distant organs. The polyclonal origins of metastasized deposits suggest that metastasis is largely driven by collective cell invasion [1,2] instead of individual cell dissemination as previously presumed. The disproportionately high metastatic potential of clusters of circulating tumor cells (CTCs) as compared to solitary CTCs, and the clinical association of CTC clusters with worse patient outcomes lends further credence to the role of collective invasion in tumor progression [3]. Thus, investigating the emergent properties enabling collective cell invasion is crucial to eventually develop anti-metastatic drugs.

Collective invasion packs often contain phenotypically distinct cell populations-‘leaders’ and ‘followers’ with varying functional properties such as remodeling of extracellular matrix (ECM), stemness, metabolism, proliferation, and immune-evasion [4–9]. This functional diversity enables co-operation among these heterogeneous populations possibly through mechano-chemical cell-cell communication, thereby amplifying the metastatic potential of such heterotypic clusters [5,10–13]. Further, the leader and follower cell-states can dynamically switch between one another, thus maintaining intratumor phenotypic heterogeneity and enriching the complexity of interactions that are responsible for aggravated metastasis driven by collectively invading cohorts of cells [6].

A key mechanism associated with collective invasion is Epithelial-Mesenchymal Transition (EMT), through which epithelial cells at least partially lose their E-cadherin mediated cell-cell adhesion and gain migratory and invasive traits and upregulate the expression of intermediate filaments such as Vimentin [14]. Initially implicitly assumed to be a binary process in cancer with epithelial (non-migratory) and mesenchymal (solitary migration) states, the concept of EMT was considered contradictory to collective cell invasion [11]. However, recent appreciation of many hybrid epithelial/mesenchymal (E/M) states with remnant cell-cell adhesion and ability to act as ‘leaders’ in collective invasion [15,16] has suggested that while cells undergoing a complete EMT may invade as single cells, partial EMT contributes to collective invasion of cells with epithelial and partial EMT states. Functional traits of cancer cells undergoing a partial EMT such as higher plasticity [17,18], tumor-initiation potential [19,20] and immune-evasion [21,22] also support the association of hybrid E/M state(s) with leader cell phenotype. Cells undergoing EMT can remodel the surrounding ECM by secreting matrix metallo-proteinases (MMPs) and enzymes such as LOXL2 that can increase ECM stiffness [23–25]. High ECM stiffness, in turn, can promote EMT, forming a mechanochemical feedback loop [26,27]. However, it remains unclear how a complex interplay among cancer cells undergoing partial or full EMT, ECM density, and stochastic invasion dynamics [28] drives the emergent outcomes of heterotypic collective cell invasion. Moreover, how each of the progressive steps in alteration of adhesion between cancer cells and their surrounding milieu affect the behavior of the population as a whole remains less understood.

Most experimental efforts investigating the role of partial or full EMT in metastasis and/or collective invasion have very limited spatiotemporal data for phenotypic heterogeneity in primary tumors [16,17]. This limitation precludes our ability to interrogate how any disruptions in spatial heterogeneity patterns or molecular perturbations in specific spatial tumor sections could impact collective invasion outcomes. Some spatially extended computational models have focused on explaining experimentally observed patterns of spatial heterogeneity in EMT status in primary tumors, but they fall short of investigating the implications of this heterogeneity in driving collective invasion [29–31]. Further, the role of heterogeneous EMT-like scenarios individually (loss of cell-cell adhesion, loss of apico-basal polarity [32]) has only begun to be appreciated in terms of diverse hybrid E/M states.

A computational model that elucidates how collective cell invasion is shaped by decrease in cell-cell adhesion and increase in cell-ECM adhesion during EMT separately, along with emergent mechanochemical dynamics of heterogeneous cell populations with each other and with ECM is, therefore, missing. Here, to overcome this limitation, we present a body of simulations wherein, within a collectively invading tumor population, a niche is allowed to undergo epithelial to mesenchymal transition with multiple alterations in adhesion tested individually or in combinations. Our results showcase how under dense ECM contexts, the intra-niche but not inter-niche adhesion strength potentiates the fitness of the transitioned niche allowing it to contribute dominantly to the collectively invading tumor.

## Results

### The niche undergoing EMT dominates tumor population at high ECM density

We started the simulations with a single cancer cell at the center of the simulation domain surrounded by ECM comprising of Collagen I fibers (Fig. 1A left). To study the influence of an epithelial to mesenchymal transition (EMT) that cancer cells undergo during the early stages of cancer metastasis and intravasation, a mesenchymal transition characterized by loss in cell-cell adhesion and gain of cell-ECM adhesion was induced in a single peripherally located cell at the 32-cell stage of tumor evolution (Fig. 1A middle). The population originating from the cell undergoing EMT shall be referred to as the transitioned population (yellow) while the parental population will be referred to as the un-transitioned population (red). The adhesion between two yellow or two red cells will be referred to as intra-niche adhesion, whereas the adhesion between a yellow and a red cell will be referred to as inter-niche adhesion. We studied how the origination of a niche characterized by EMT influences the various phenotypic traits of the tumor undergoing collective cell invasion (CCI) which will be assessed as before [33] by measuring the area of largest connected cluster of cells (Fig. S1). We introduced another output for this study namely, cell ratio, which is the ratio of ‘transitioned to ‘un-transitioned’ population numbers that quantifies the impact and fitness of the individual niches and when ratioed as above, quantifies the domination of the transitioned niche. Along with these, we track two other intermediate quantities: cumulative levels of matrix metalloproteinases (MMP) secreted from all cancer cells of both types and cumulative levels of growth factor (GF) chemical released from the ECM fibers, both which influence the tumor evolution.

**FIGURE 1:**
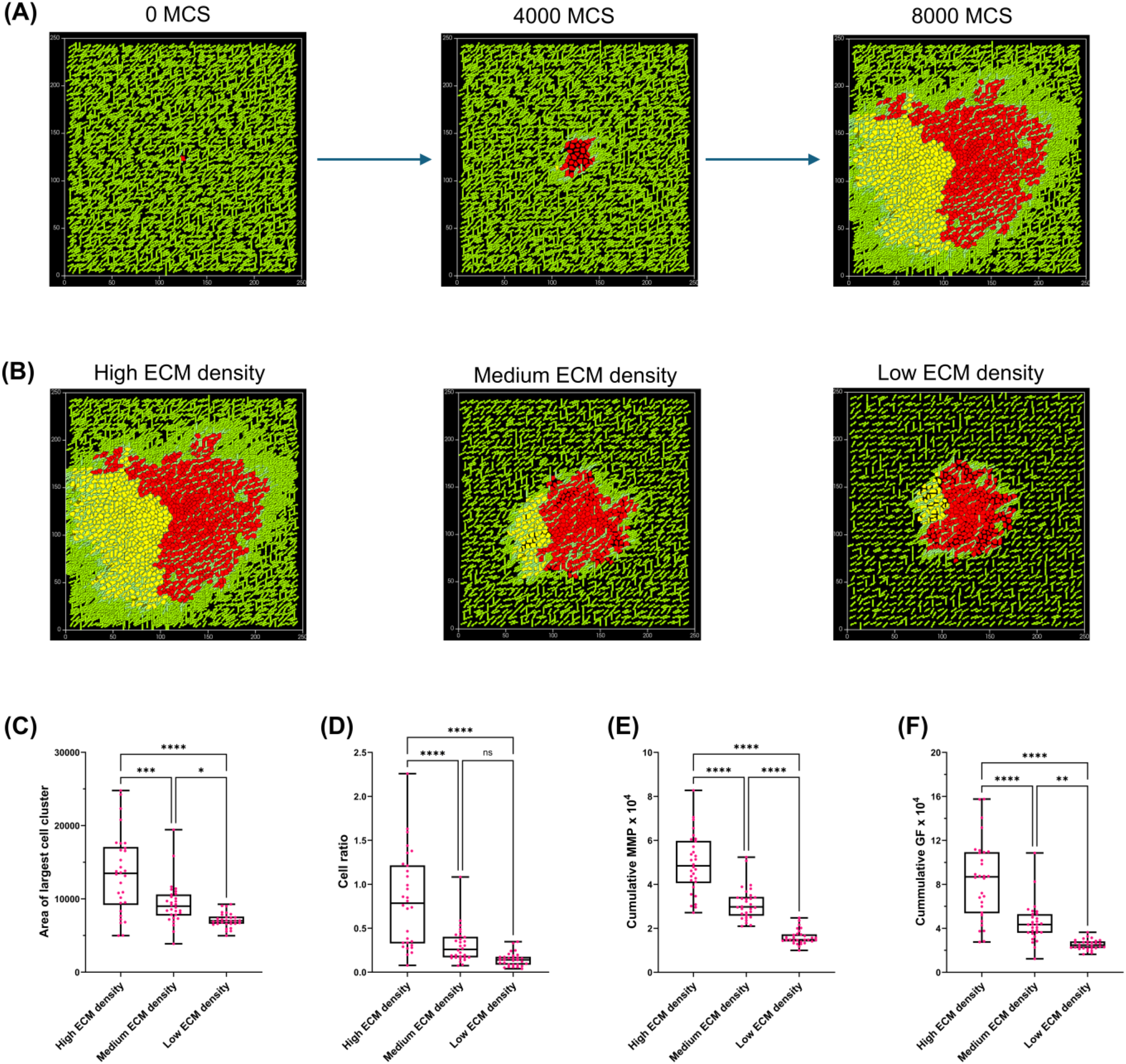
Spatiotemporal collective dynamics of a cellular niche showing epithelial to mesenchymal transition (EMT) in competition with its un-transitioned parent population. (A) Starting arrangement of the simulation with a single cancer cell (red) at the center of the simulation domain surrounded by extracellular matrix (ECM) fibers (left), undergoing collective invasion with an introduction of a single cell (yellow; center) at the 32 cell stage of tumor evolution showing an epithelial to mesenchymal transition characterized by low inter- and intra-niche adhesion and high cell-ECM adhesion. Final state of the tumor (8000 MCS) consisting of competing populations of the transitioned niche (yellow) and the un-transitioned niche (red). (B) Tumor population phenotypes at the end point of simulations (8000 MCS) with interspacing between ECM fibers decreasing from left to right (i.e., high ECM density on the left, medium ECM density in the center and low ECM density on the right) (C) Box and whisker plot with median values of the area of largest cell cluster (used as a measure of collective cancer invasion) for 30 replicates of simulations for high, medium, and low ECM densities. (D) Box and whisker plot with median values of the cell ratio (used as a measure of impact of the EMT of one niche on overall tumor composition and invasion) for 30 replicates of simulations for high, medium, and low ECM densities. (E) Box and whisker plot with median values of the cumulative levels of matrix metalloproteinases (MMP) secretions from all cancer cells for 30 replicates of simulations for high, medium, and low ECM densities. (F) Box and whisker plot with median values of the cumulative levels of growth factor (GF) chemical secretions from ECM fibers for 30 replicates of simulations for high, medium, and low ECM densities. Statistical significance for all measurements in the four graphs was computed using one-way ANOVA with Tukey’s post hoc multiple comparisons. Significance (p-value) is represented as *, where * implies p≤0.05, ** implies p ≤0.01, *** implies p ≤ 0.001, and **** implies p ≤ 0.0001.

Our previous work had identified adhesion strengths at which collective cancer invasion (CCI) was observed [33] and these cell-cell and cell-ECM adhesions strengths (See Supplementary Information for parameter values) were assigned to the un-transitioned parental population cells. To begin with, we induce a complete EMT in a peripherally located cell at the 32 cell-stage of the tumor by assigning inter- and intra-cell adhesion strength for transitioned cells (yellow) at half of that of the un-transitioned cells and the cell-ECM adhesion strength at double that of un-transitioned cells. These simulations were carried out in three different ECM densities that are obtained by gradually increasing the interspacing distance between ECM fibers (d) from 3 to 5 voxels (Fig. 1B). The ECM with the least interspacing is considered to have high ECM density, while the one with the highest interspacing is representative of low ECM density and the intermediate voxel spacing represented medium ECM density.

We observed that Collective cancer invasion (CCI) decreased as the ECM became sparse (Fig. 1C, ANOVA p<0.0001; significance assessed by Tukey’s post hoc multiple comparisons). The domination of transitioned cell population was evident at high ECM density conditions (Fig. 1D, ANOVA p<0.0001; differences between cell ratios in medium and low-density cases were significant compared to high density cases as assessed by Tukey’s post hoc multiple comparisons). Cumulative MMP and GF secretion levels progressively decreased with decrease in ECM density, implying reduced chemical activity as the ECM became sparser (Fig. 1E and 1F respectively, ANOVA p<0.0001; differences between levels for both under different density conditions significant as assessed by Tukey’s post hoc multiple comparisons).

In simulations under these conditions, across multiple replicates, we observed a variation in the end phenotype: in some cases, transitioned cells were surrounded by cells of the parental population (un-transitioned cells) and showed less invasion, whereas in most simulations, the transitioned population were not surrounded by the former and showed greater invasion (Fig. 2A left and right respectively showing simulation micrographs at 8000 MCS of two end phenotypes). We chose a threshold value for cell ratio of 0.3 to classify the end phenotypes of cell populations into the above two outcomes: if the ratio of the end point phenotype of tumor was less than 0.3 then we classified that case as ‘confined’, otherwise it was classified as ‘unconfined’ (Fig. 2B shows a pie chart depicting probability of occurrence of confined being 18% while that of unconfined being 82%). The unconfined simulations showed higher median value of CCI than control cases of ‘no transition’, whereas the CCI for confined case simulations was insignificant (Fig. 2C; ANOVA p<0.0001; significance as assessed by Tukey’s post hoc multiple comparisons). EMT significantly increased the domination of the transitioned niche, when simulations (60 replicates) were stratified based on confinement (Fig. 2D; ANOVA p<0.0001; significance as assessed by Tukey’s post hoc multiple comparisons). The cumulative levels of MMPs and GFs for unconfined cases were significantly higher than control levels (Fig. 2E and F; ANOVA<0.0001 significance as assessed by Tukey’s post hoc multiple comparisons). These observations highlight the impact of confinement effect and EMT on inter-niche competition and dynamics.

**FIGURE 2:**
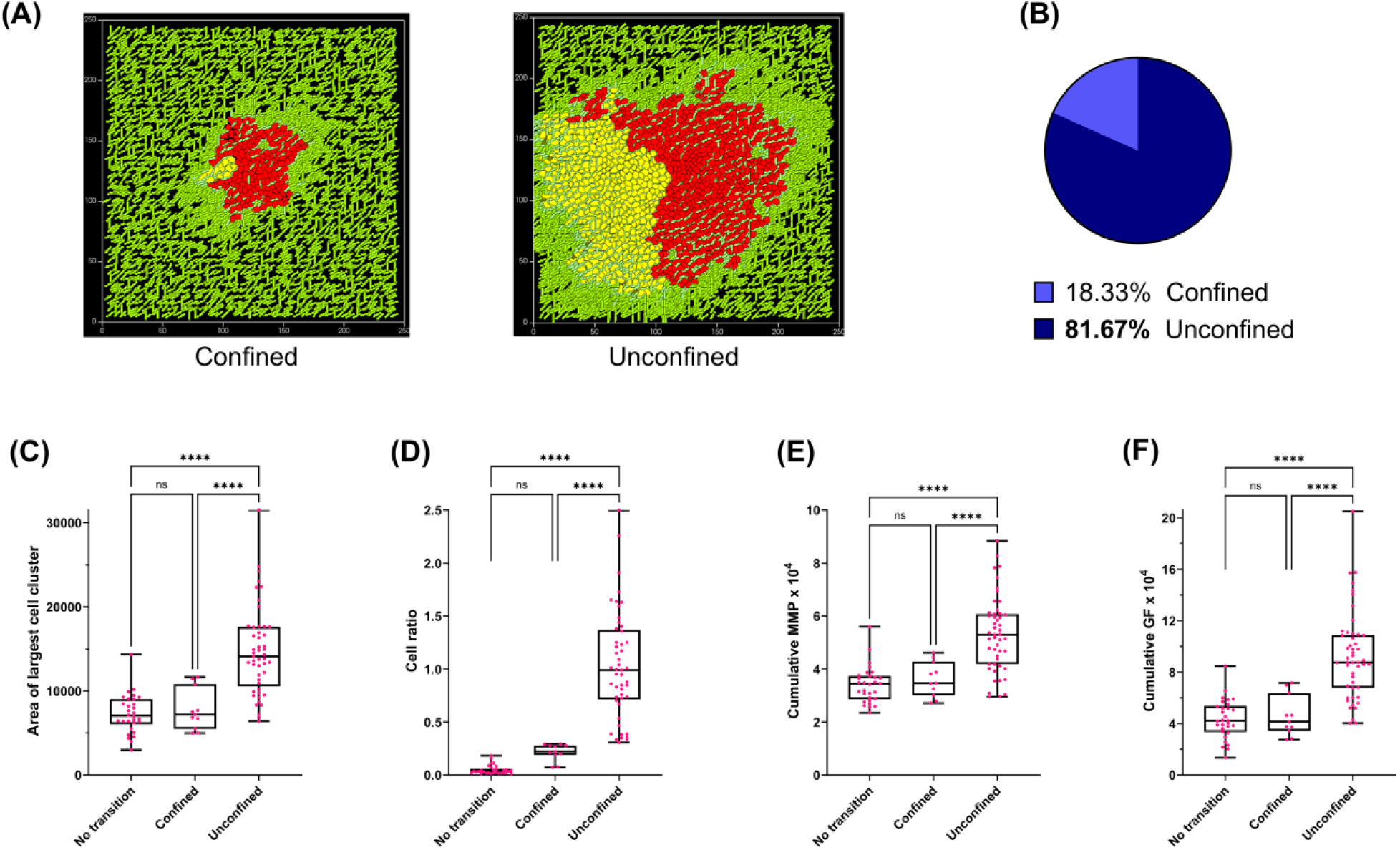
Complete Epithelial to Mesenchymal Transition (EMT) of transitioned niche shows domination over the un-transitioned one with 82% probability in a high ECM density tumor microenvironment. (A) Image showing transitioned cells (yellow) confined by un-transitioned parent population (red) (left image), transitioned cells (yellow) unconfined by un-transitioned parent population (red) (right image) at the end point of simulations (8000 MCS). (B) Pie chart showing the probability of occurrence of confined (light blue) and unconfined (dark blue) cases. (C) Box and whisker plot with medians for area of largest cell cluster (used as a measure of collective cancer invasion) for the control case (no transition), for confined and unconfined cases of complete EMT. (D) Box and whisker plot with medians for the cell ratio (used as measure of impact of transitioned niche) for the control case (no transition), for confined, and unconfined cases of complete EMT. (E) Box and whisker plot with medians of cumulative levels of matrix metalloproteinases (MMP) secretions from all cancer cells, for the control case (no transition), for confined, and unconfined cases of complete EMT. (F) Box and whisker plot with medians of cumulative levels of growth factor (GF) secretions from the ECM fibers, for the control case (no transition), for confined, and unconfined cases of complete EMT. Statistical significance for all measurements in the four graphs was computed using one-way ANOVA with Tukey’s post hoc multiple comparisons. Significance (p-value) is represented as *, where * implies p≤0.05, ** implies p ≤0.01, *** implies p ≤ 0.001, and **** implies p ≤ 0.0001.

### Increasing cell-ECM adhesion and decreasing cell-cell adhesion have distinct consequences

EMT could be construed as a multistep process with at least three discrete events: an increase in cell-ECM adhesion, a decrease in adhesion between the transitioning and parental niche (inter-niche adhesion) as well as among cells of the transitioning niche (intra-niche adhesion). To study how each of these events individually contribute to CCI and transitioned niche domination, we implemented distinct sets of EMT events in the dense ECM milieu. The first set is a gain in cell-ECM adhesion in the transitioned cells, compared with the parental cell population at the time of induction of EMT in the tumor at a 32-cell stage. Both the above-characterized end phenotypes were observed for these simulations (Fig. 3A left and right showing simulation micrographs at 8000 MCS for confined and unconfined cases, respectively; see Videos S1 and S2). We observed that increased cell-ECM adhesion transition was able to achieve domination only in 50% of replicates (30 out of 60 simulation replicates were unconfined; Fig. 3B shows a pie chart depicting probability of occurrence of unconfined and confined as end phenotypes). Similar to the complete EMT cases, the unconfined simulations showed higher CCI than control cases of no transition, whereas the CCI for confined simulations was insignificantly different (Fig. 3C; ANOVA p<0.0001; significance as assessed by Tukey’s post hoc multiple comparisons). Increase in cell-ECM adhesion transition increased the domination of the unconfined transitioned niche simulations (Fig. 3D; ANOVA p<0.0001; significance as assessed by Tukey’s post hoc multiple comparisons). The cumulative levels of MMPs and GFs for unconfined cases were higher than control levels (Fig. 3E and F; ANOVA<0.0001; significance as assessed by Tukey’s post hoc multiple comparisons). For all the metrics: CCI, niche domination and their molecular counterparts the median values for the unconfined cases with only cell-ECM adhesion increase were lesser than the complete EMT cases.

**FIGURE 3:**
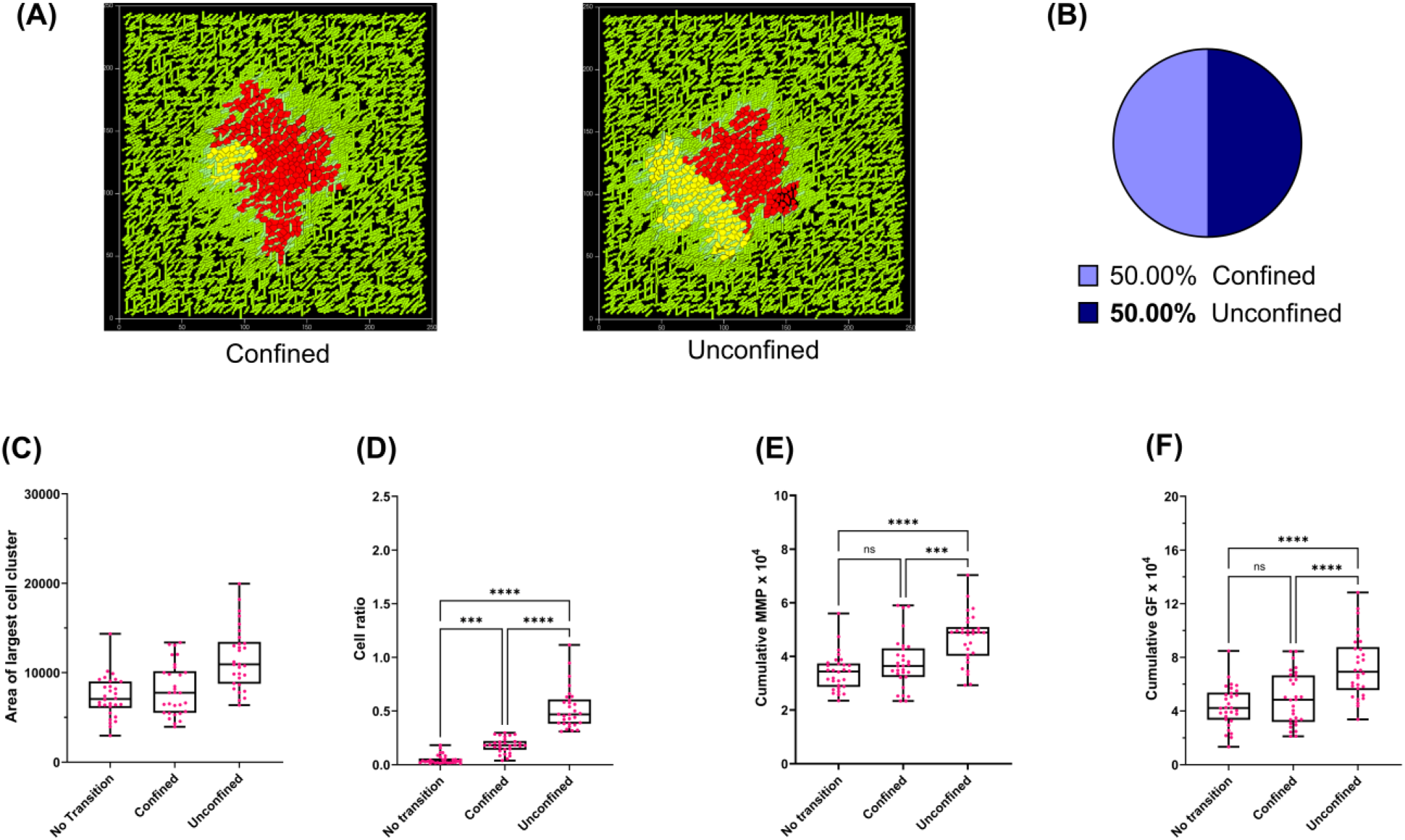
Increased cell-ECM adhesion allows transitioned niche domination with a 50% probability. (A) Image showing transitioned niche (yellow) confined by un-transitioned niche (red) (left image), and transitioned niche unconfined by un-transitioned niche (right) at the end point of simulation (8000 MCS). (B) Pie chart showing the probability of occurrence of confined (light blue) and unconfined (dark blue) cases. (C) Box and whisker plot with medians for area of largest cell cluster (used as a measure of collective cancer invasion) for the control case (no transition), and for simulations wherein the transitioned cells have higher cell-ECM adhesion for confined and unconfined cases. (D) Box and whisker plot with medians for the cell ratio (used as measure of impact of transitioned niche) for the control case (no transition), and for simulations wherein the transitioned cells have higher cell-ECM adhesion for confined and unconfined cases. (E) Box and whisker plot with medians of cumulative levels of matrix metalloproteinases (MMP) secretions from all cancer cells, for the control case (no transition), and for simulations wherein the transitioned cells have higher cell-ECM adhesion for confined and unconfined cases. (F) Box and whisker plot with medians of cumulative levels of growth factor (GF) secretions from the ECM fibers, for the control case (no transition), and for simulations wherein the transitioned cells have higher cell-ECM adhesion for confined and unconfined cases. Statistical significance for all measurements in the four graphs was computed using one-way ANOVA with Tukey’s post hoc multiple comparisons. Significance (p-value) is represented as *, where * implies p≤0.05, ** implies p ≤0.01, *** implies p ≤ 0.001, and **** implies p ≤ 0.0001.

In the second set, we simulated a reduction in both inter- and intra-niche cell-cell adhesion with no change in cell-ECM adhesion, within 32-cell tumor under high ECM density. The probability of such transitioned niches showing domination was only 15% (9 out of 60 simulation replicates were unconfined; Fig. 4A left and right showing simulation micrographs at 8000 MCS for confined and unconfined cases; Fig. 4B shows a pie chart depicting probability of domination by the transitioned niche). The unconfined simulation cases showed a significant increase in CCI, although the median value was lower than for complete EMT (Fig. 2; Fig. 4C; ANOVA p<0.0001 significance as assessed by Tukey’s post hoc multiple comparisons). The unconfined transitioned niche showed a significantly higher cell ratio compared with controls, but the median value was lower than for the unconfined counterparts with full EMT and close to the unconfined cases of earlier high cell-ECM adhesion transition (Fig. 4D; ANOVA<0.0001 significance as assessed by Tukey’s post hoc multiple comparisons). The unconfined simulations exhibited significantly higher MMP and GF levels (with mild differences with the equivalent cases representing cell-ECM adhesion but lower than the complete EMT unconfined cases), the same for the majority of simulations, i.e., confined cases were insignificantly different from controls (Fig. 4E and F; ANOVA<0.0001 significance as assessed by Tukey’s post hoc multiple comparisons). The drastic reduction of probability of domination by the transitioned niche from 50% to 15% thus indicated that more than loss of inter- and intra-niche adhesions, gain of cell-ECM adhesion relieved, albeit partially and occasionally, the confinement of mesenchymally transitioned niche by surrounding un-transitioned cells.

**FIGURE 4:**
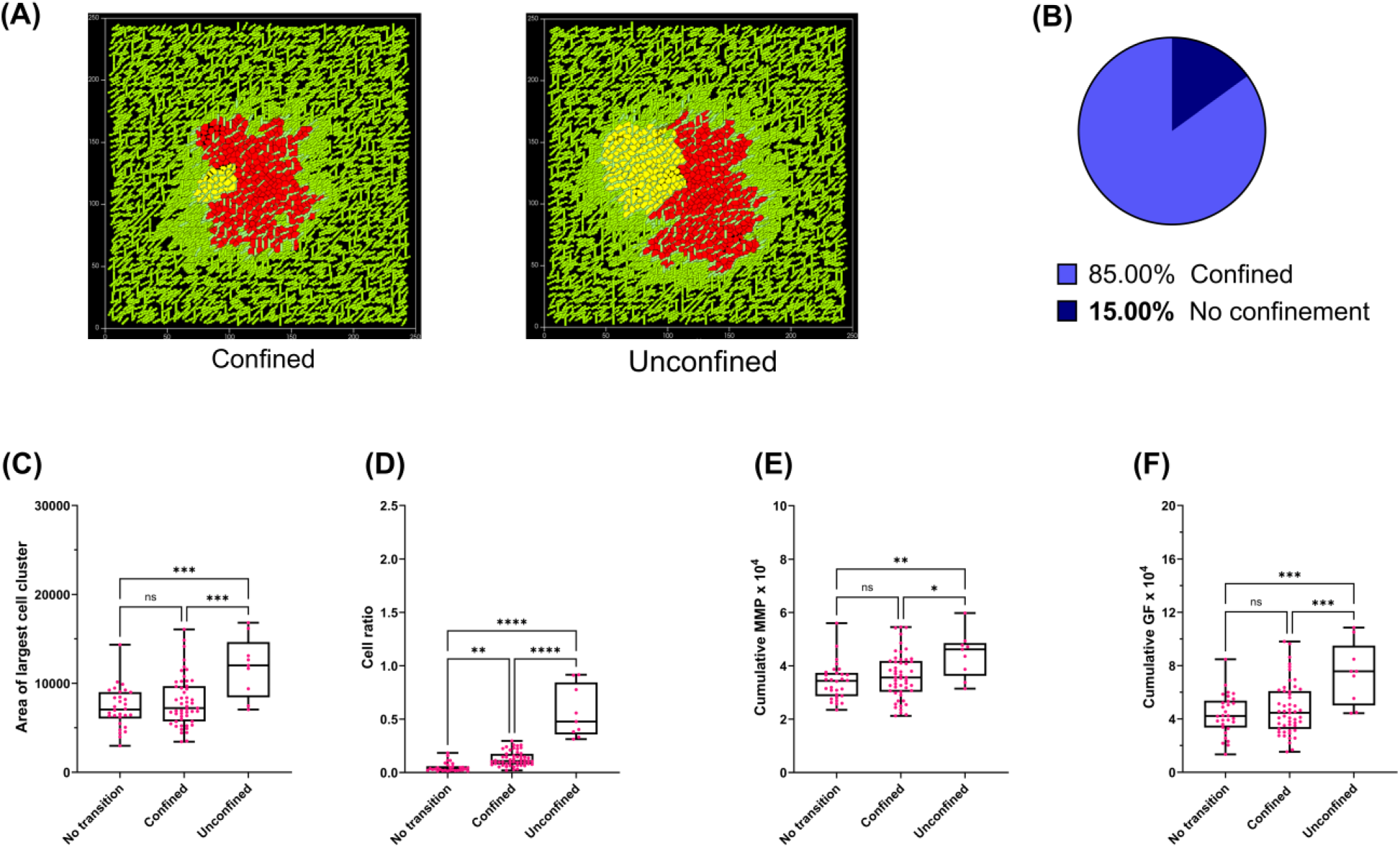
Reduced inter- and intra-cell adhesions do not allow transitioned niche domination. A) Image showing transitioned cells (yellow) confined by un-transitioned niche (red) (left image), transitioned cells unconfined by un-transitioned niche (right) at the end point of simulation (8000 MCS). (B) Pie chart showing the probability of occurrence of confined (light blue) and unconfined (dark blue) cases. (C) Box and whisker plot with medians for area of largest cell cluster (used as a measure of collective cancer invasion) for the control case (no transition), and for simulations, wherein the transitioned cells have lower intra- and inter-niche adhesion for confined and unconfined cases. (D) Box and whisker plot with medians for the cell ratio (used as measure of impact of transitioned niche) for the control case (no transition), and for simulations wherein the transitioned cells have lower intra- and inter-niche adhesion for confined and unconfined cases. (E) Box and whisker plot with medians of cumulative levels of matrix metalloproteinases (MMP) secretions from all cancer cells, for the control case (no transition), and for simulations wherein the transitioned cells have lower intra- and inter-niche adhesion for confined and unconfined cases. (F) Box and whisker plot with medians of cumulative levels of growth factor (GF) secretions from the ECM fibers, for the control case (no transition), and for simulations wherein the transitioned cells have lower intra- and inter-niche adhesion for confined and unconfined cases. Statistical significance for all measurements in the four graphs was computed using one-way ANOVA with Tukey’s post hoc multiple comparisons. Significance (p-value) is represented as *, where * implies p≤0.05, ** implies p ≤0.01, *** implies p ≤ 0.001, and **** implies p ≤ 0.0001.

### Loss of Intra-niche and inter-niche adhesion have different impacts on niche dominance

We next asked if individually implementing a decrease in cell adhesion within the transitioned niche (intra-niche adhesion) or between the transitioned niche and the parental population (inter-niche adhesion) in the background of higher cell-ECM adhesion transition was sufficient to evince the complete phenotypic behavior of a niche with complete EMT effect as seen in Figure 2. Combining low inter-niche cell adhesion with high cell-ECM adhesion transition achieved unconfined transitioned niches in 48% simulations which is almost same as that of mere high cell-ECM adhesion transition (Fig. 5A left and right showing simulation micrographs at 8000 MCS for confined and unconfined cases; Fig. 5B shows a pie chart depicting probability of domination by the transitioned niche). Such unconfined simulations exhibited higher CCI (Fig. 5C; ANOVA<0.0001 significance as assessed by Tukey’s post hoc multiple comparisons), showed greater niche domination (Fig. 5D; ANOVA<0.0001 significance as assessed by Tukey’s post hoc comparison) and also showed significantly higher levels of MMP and GF secretions (Fig. 5E and F; ANOVA<0.0001 significance as assessed by Tukey’s post hoc multiple comparisons). Unsurprisingly, the median values of all the above-mentioned metrics were lower than for the complete EMT cases.

**FIGURE 5:**
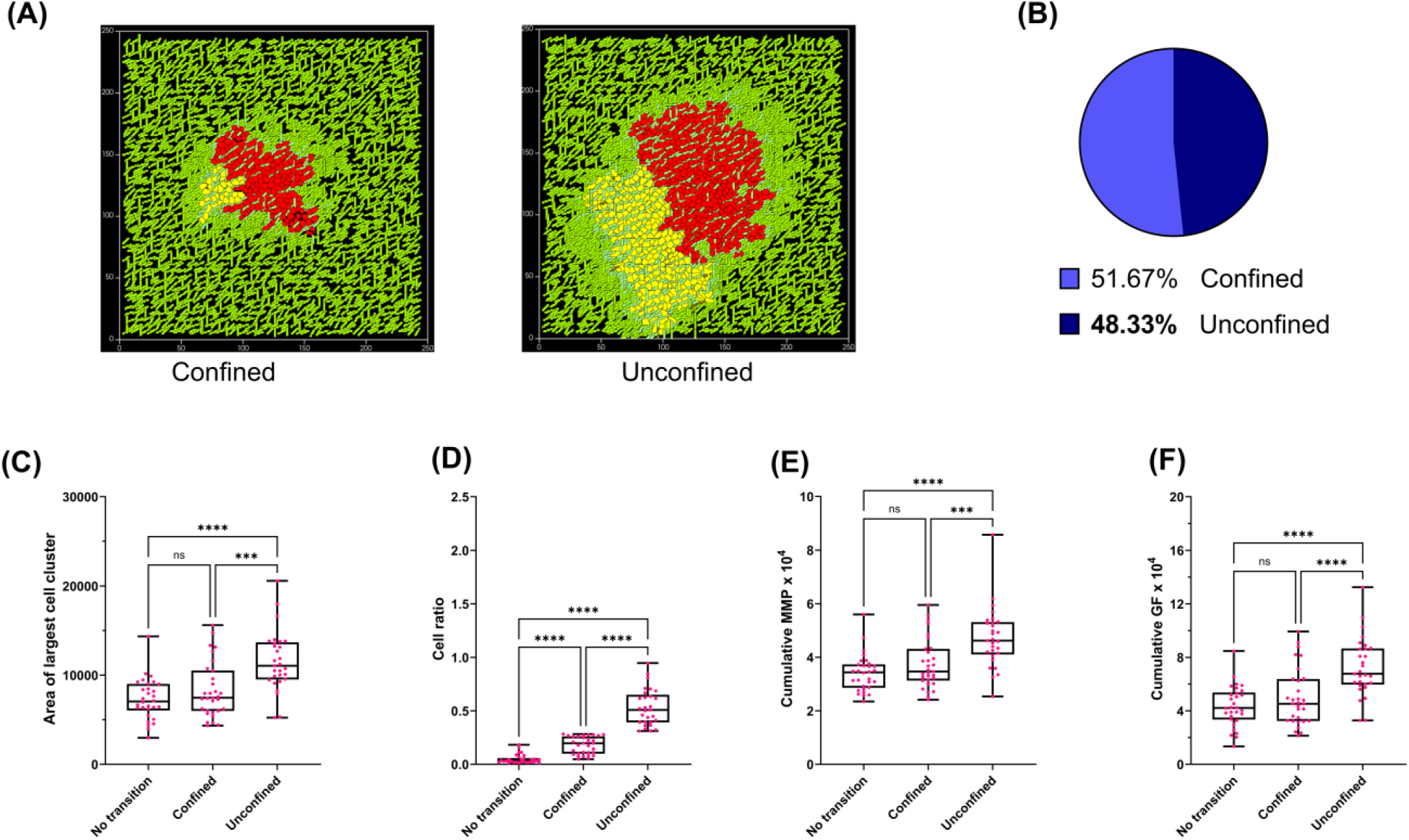
Probability of transitioned niche domination is unaffected by addition of reduced inter-cell adhesion to high cell-ECM adhesion. (A) Image showing transitioned cells (yellow) confined by un-transitioned niche (red) (left image), transitioned cells unconfined by un-transitioned niche (right) at the end point of simulation (8000 MCS). (B) Pie chart showing the frequency/probability of occurrence of confined (light blue) and unconfined (dark blue) cases. (C) Box and whisker plot with medians for area of largest cell cluster (used as a measure of collective cancer invasion) for the control case (no transition), and for simulations wherein the transitioned cells have higher cell-ECM adhesion and lower inter-niche adhesion for confined and unconfined cases. (D) Box and whisker plot with medians for the cell ratio (used as measure of impact of transitioned niche) for the control case (no transition), and for simulations wherein the transitioned cells have higher cell-ECM adhesion and lower inter-niche adhesion for confined and unconfined cases. (E) Box and whisker plot with medians of cumulative levels of matrix metalloproteinases (MMP) secretions from all cancer cells, for the control case (no transition), and for simulations wherein the transitioned cells have higher cell-ECM adhesion and lower inter-niche adhesion for confined and unconfined cases. (F) Box and whisker plot with medians of cumulative levels of growth factor (GF) secretions from the ECM fibers, for the control case (no transition), and for simulations wherein the transitioned cells have higher cell-ECM adhesion and lower inter-niche adhesion for confined and unconfined cases. Statistical significance for all measurements in the four graphs was computed using one-way ANOVA with Tukey’s post hoc multiple comparisons. Significance (p-value) is represented as *, where * implies p≤0.05, ** implies p ≤0.01, *** implies p ≤ 0.001, and **** implies p ≤ 0.0001.

In contrast to the above-described observations, implementing a loss in intra-niche adhesion (among the cells of the transitioned niche) in the background of increased cell-ECM adhesion resulted in recovery of niche domination probability (82%) equivalent to the complete EMT (Fig. 5A left and right showing simulation micrographs at 8000 MCS for confined and unconfined cases; Fig. 5B shows a pie chart depicting probability of occurrence of unconfined and confined cases). Unconfined cases showed higher CCI than controls (although median value was lower than not just the complete EMT set but even the unconfined set with just an increase in cell-ECM adhesion; Fig. 6C; ANOVA<0.0001 significance as assessed by Tukey’s post hoc multiple comparisons). Cell ratio for unconfined simulations is higher than controls (with a median value higher than for the unconfined simulations with high cell-ECM adhesion but lower than for the complete EMT set), (Fig. 6D; ANOVA<0.0001 significance as assessed by Tukey’s post hoc multiple comparisons). MMP and GF levels for the unconfined cases were significantly higher than controls (with mild differences with the equivalent cases representing cell-ECM adhesion but lower than the complete EMT unconfined cases) (Fig. 6E and F; ANOVA<0.0001 significance as assessed by Tukey’s post hoc multiple comparisons). Taken together, our results indicate that the inter-niche adhesion operating between the transitioned and un-transitioned niches contribute together to the overall tumor evolution, the probability of domination by the transitioned niche is predominantly driven within the niche through the adhesion of its cells (intra-niche adhesion) to each other and to the ECM.

**FIGURE 6:**
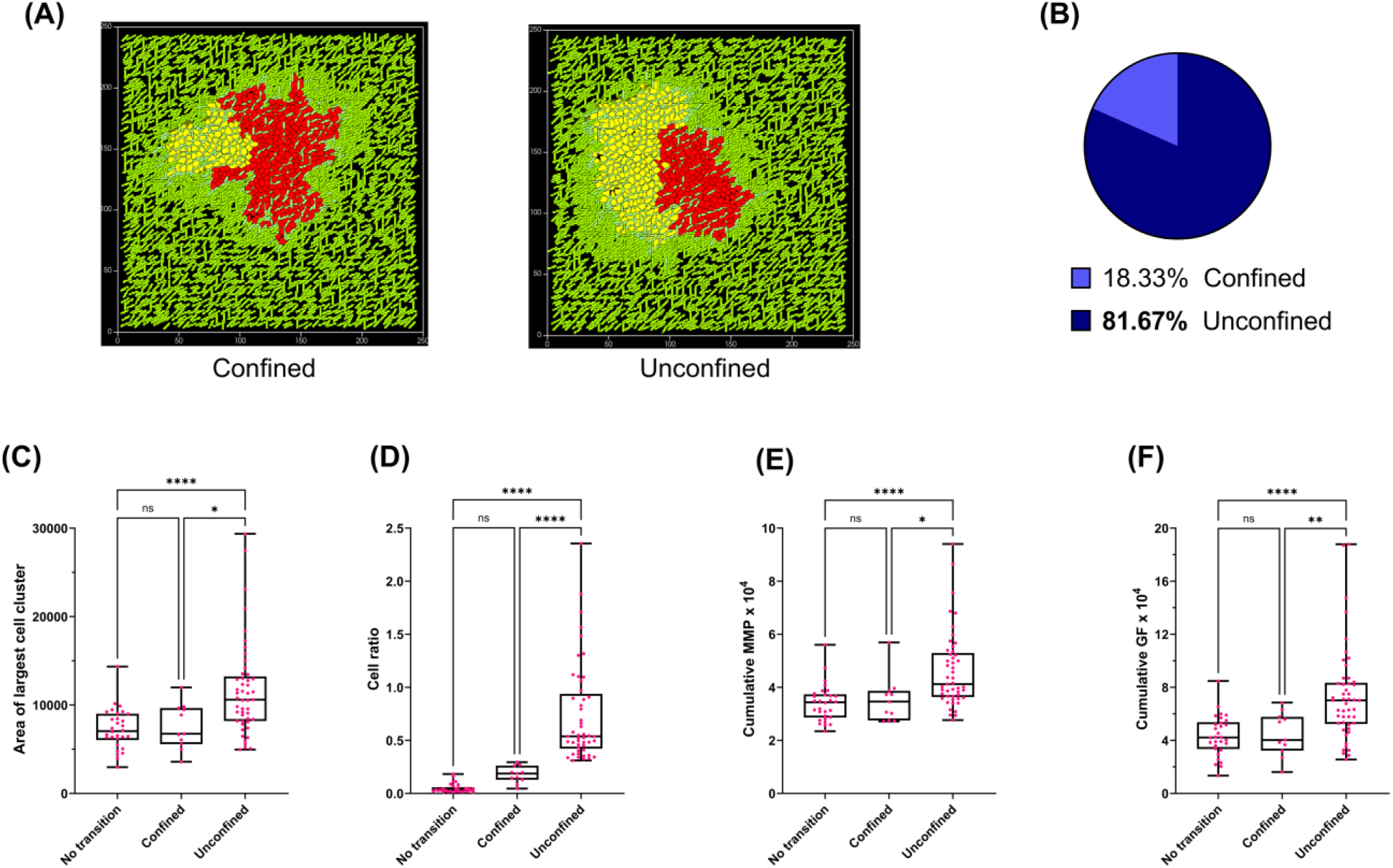
Higher cell-ECM adhesion combined with lower intra-niche adhesion is sufficient for the transitioned niche to show domination. (A) Image showing transitioned cells (yellow) confined by un-transitioned niche (red) (left image), transitioned cells unconfined by un-transitioned niche (right) at the end point of simulation (8000 MCS). (B) Pie chart showing the probability of occurrence of confined (light blue) and unconfined (dark blue) cases. (C) Box and whisker plot with medians for area of largest cell cluster (used as a measure of collective cancer invasion) for the control case (no transition), and for simulations wherein the transitioned cells have higher cell-ECM adhesion and lower intra-niche adhesion for confined and unconfined cases. (D) Box and whisker plot with medians for the cell ratio (used as measure of impact of transitioned niche) for the control case (no transition), and for simulations wherein the transitioned cells have higher cell-ECM adhesion and lower intra-niche adhesion for confined and unconfined cases. (E) Box and whisker plot with medians of cumulative levels of matrix metalloproteinases (MMP) secretions from all cancer cells, for the control case (no transition), and for simulations wherein the transitioned cells have higher cell-ECM adhesion and lower intra-niche adhesion for confined and unconfined cases. (F) Box and whisker plot with medians of cumulative levels of growth factor (GF) secretions from the ECM fibers, for the control case (no transition), and for simulations wherein the transitioned cells have higher cell-ECM adhesion and lower intra-niche adhesion for confined and unconfined cases. Statistical significance for all measurements in the four graphs was computed using one-way ANOVA with Tukey’s post hoc multiple comparisons. Significance (p-value) is represented as *, where * implies p≤0.05, ** implies p ≤0.01, *** implies p ≤ 0.001, and **** implies p ≤ 0.0001.

Our results suggest that higher cell-ECM adhesion and lower intra-niche cellular adhesion combinedly ensure as high a probability of niche domination as the complete EMT within high density ECM milieu.

## Discussion

The cues that initiate EMT can be non-cell-autonomous (such as increased TGF-β levels, decreased oxygen, increased elastic moduli of substrata, increased levels of specific ECM proteins such as fibronectin [34–38]) or autonomous (mutations or epigenetic perturbations in signaling hub-proteins such as EGFR, Ras, Src, FAT1, p53 etc [39–44]). Heterogeneity in the presentation and pervasion of extracellular cues or occurrence of secondary mutations entails that EMT may be niche specific [45,46]. Given the intense competition for survival between distinct niches within a growing tumor, it is pertinent to ask how the altered phenotypic state of an EMT niche regulates its fitness vis-a-vis its parental un-transitioned population.

It was to answer this question that we ‘induced’ an EMT event in a single cell within a growing 32-cell tumor in our computational model and traced the evolution of both the parental and transitioned lineages as a function of their intrinsic behaviors, their interactions with each other and the ECM microenvironment. In the control case of ‘No transition’ (where both these niches are distinct only through their color and not any other assigned properties), the new lineage was almost invariably confined by the original cellular population, due to an asymmetric beginning cell number. This setup was appropriate to deploy in asking what properties of EMT under what microenvironmental contexts enable a fledgling niche to survive such a confinement and in fact begin to even dominate the parental population. Our simulations reveal high cell-ECM adhesion combined with low intra-niche adhesion being pivotal in complete EM Transitioned niche to achieve domination.

The first important criterion for the domination by a completely EM transitioned niche, that we observed was the density of the surrounding stromal Collagen I-like fibrillar ECM. At progressively lower densities of ECM, collective migration is impeded due to lower rates of attachment and detachment to ECM fibers by migrating cells [47,48]. Moreover, optimal densities of ECM represent the ‘sweet spot’ wherein migrating masses are sufficiently jammed such that they can collectively migrate even if intra-niche adhesion is low [33,49]. Interestingly, stochasticity plays an important role in determining the ability of each step of EMT to contribute to higher fitness of its cellular niche. Invoking only an increased cell-ECM adhesion (the mesenchymal arm of EMT) achieves domination in half the simulations. Although this is a function of the ‘time’ at which the EMT is introduced in the simulations, this means the later the introduction, the lower the probability with which accentuating the cell-ECM adhesion solely will allow the niche to outcompete its parental cells. The stochastic regulation of domination probability can be traced to the geometry of the interface between the two populations: not just should the transition originate at the edge of a cell population the subsequent population dynamics should ensure that the transitioned cell is not quickly surrounded and confined by un-transitioned parental cells. We observed that decreasing adhesion among transitioned cells (intra-niche) rather than decreasing adhesion between the transitioned and un-transitioned cells (inter-niche) (in addition to an increase in cell ECM adhesion) increased the probability of the EMT-driven niche domination. This could be because as the niche grew, the number of weakened cell-cell adhesion junctions contributing to the phenotype increased for the former case much more than the latter.

Careful observations of the chemical fields reveal another factor at play: when cell-ECM adhesion is enhanced, the transitioned cells degrade the ECM better releasing factors that serve to increase growth and chemoattract cells [50–53]. This effect is not limited just to transitioned cells but also to un-transitioned cells which outnumber the former in the initial stages, chemotaxis towards growth factor concentration near transitioned niche influenced by ECM fibers orientation and contribute to confinement of transitioned niche (see Video V1, V2). The rapidly surrounding un-transitioned cells and the transitioned ones also produce fresh ECM which further accentuates the confining effect by reducing the contact between transitioned cells with existing Collagen I fibers. Decreasing the intra-niche adhesion competes with this confining effect allowing the transitioned niche to invade faster.

We would like to conclude by highlighting the limitations of our study. Here we have focused exclusively on the input metric combinations (of adhesion, ECM secretion and degradation etc), which are cognate to collective cell invasion. We have not focused on dispersed modes of migration wherein a separate set of rules may determine mesenchymal domination. Computational power (especially associated with the construction of the ECM) limits our simulations to be two-dimensional. In addition, this study does not explore the role of other geometric parameters of the ECM, such as cross linking, (an)isotropic distribution, etc on EMT niche fitness. Such explorations in 3D would be undertaken in subsequent studies. These limitations do not take away from the importance of our observation that the domination of a mesenchymally transitioned niche is driven by cell autonomous (adhesive) and cell-non-autonomous (ECM density) cues.

## Methods

### Modelling framework and CompuCell3D

CompuCell3D is an agent-based simulation framework that is developed to computationally model systems in cell-, developmental- and cancer biology. It is primarily based on the Cellular Potts model (CPM)/Glazier-Graner-Hogeweg (GGH) model [54]. CompuCell3D (CC3D) enables us to combine CPM/GGH with partial differential equation (PDE) solvers for chemical fields and other models for spatiotemporal modeling by defining spatially extended generalized cells, which can represent single biological cells, their clusters or even sub-compartments and subdomains of non-cellular materials [55]. Each voxel in the simulation domain is represented by its position vector 𝒙 and a collection of a few such voxels constitutes a generalized cell. The generalized cells have a unique index 𝜎(𝒙) like cell number (𝜎 = 1,2. . 𝑁) and all the voxels inside a given generalized cell share a common index 𝜎. Each of these generalized cells are associated with a generalized cell type 𝜏. The time evolution of simulations in CC3D involves randomly selecting a lattice site 𝒙 (target voxel) and selecting another voxel in its neighborhood 𝒙′ (source voxel) followed by making an attempt to copy the cell index 𝜎(𝒙′) present at 𝒙′ on to 𝜎(𝒙) present at 𝒙 given that source and target voxel doesn’t correspond to the same generalized cell. The energy change (Δ𝐻) due to this proposed change in domain configuration is evaluated and the probability of successful voxel copying attempt is given the Boltzmann acceptance function shown below. Every Monte Carlo Step (MCS) involves multiple such voxel copying attempts, where modified Metropolis algorithm is used to spatially sample the target voxels in the simulation domain.

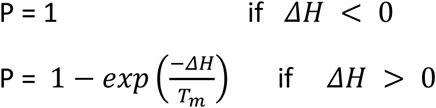

Where ΔH is change in effective energy if the copy occurs and 𝑇_𝑚_ is a parameter describing the amplitude of cell-membrane fluctuations [55].

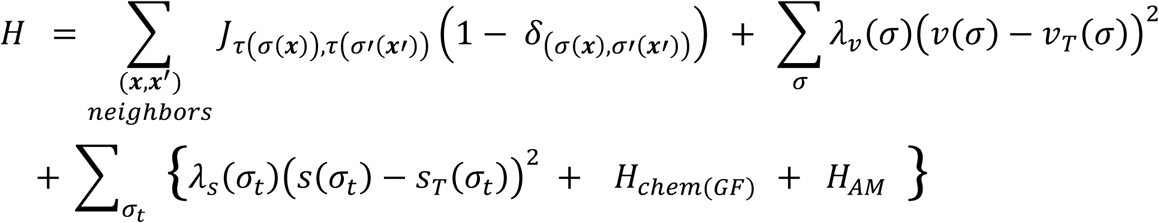

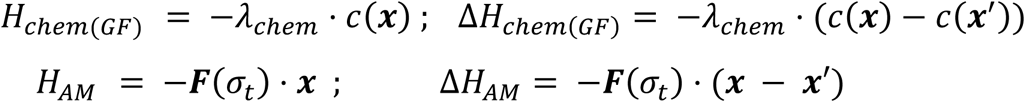

The CPM effective energy (𝐻), is the cornerstone of all CC3D simulations that describe cell properties, their interactions, and behaviors. The first term in (𝐻) is called the adhesion or boundary energy term that determines the nature and extent of interaction between various cell types. 𝐽_𝑖,𝑗_ is the contact energy between two cell types i, j and is an element of the symmetric matrix 𝐽. Contact energy is inversely related to adhesion strength, low values of contact energy imply high adhesion between/ within the cell types and vice versa. (1 − 𝛿_(𝜎(𝒙),𝜎′(𝒙′))_) term ensures that the voxel 𝒙 and its neighbor 𝒙′ doesn’t belong to the same generalized cell, thus accounting only for contact energies between boundary voxels of cells (variables in bold represent vectors in equations). The second term in effective energy is the volume constraint energy term that controls the cell size / volume by ensuring the cell volume, 𝑣(𝜎) does not deviate much from the cell’s target volume 𝑣_𝑇_(𝜎). 𝜆_𝑣_(𝜎) represents stiffness of the cell (or) inverse of compressibility of the cell. The third term in effective energy is the surface constraint energy term that influences cell shape by ensuring the cell surface, 𝑠(𝜎) does not deviate much from the cell’s target surface 𝑠_𝑇_(𝜎). 𝜆_𝑠_(𝜎) is equivalent to stiffness of cell membrane. The surface constraint term, 𝐻_𝑐ℎ𝑒𝑚(𝐺𝐹)_ and 𝐻_𝐴𝑀_terms of the effective energy (𝐻) are specific to cancer cells only in the simulations. Cell index 𝜎 corresponds to all the cells (cancer cells in tumor, existing and new ECM fibers, …) in the simulation domain while 𝜎_𝑡_ has cell indices of only cancer cells that constitute the tumor as indicated in the effective energy equation. Δ𝐻_𝑐ℎ𝑒𝑚(𝐺𝐹)_ enables chemotaxis of cancer cells that represent a biased motility of cells in response to the growth factor chemical gradient they are exposed to. 𝑐(𝒙) represents the concentration of the chemical field (GF) at the index-copying target voxel (𝒙) while 𝑐(𝒙′) corresponds to the same at the source voxel (𝒙′). 𝜆_𝑐ℎ𝑒𝑚_ is the strength of the chemotaxis that cells respond to. 𝛥𝐻_𝐴𝑀_is used to model the forces between the cells that contribute to their Brownian motion (AM in the subscript corresponds to active motility of the cells). (𝒙 − 𝒙′) is the difference between position vectors of target and source voxels. 𝑭(𝜎_𝑡_) is the force acting on cells that has both x and y components. Both F_x_, F_y_ are sampled from a uniform random distribution between (−10,10) that varies the magnitude of forces along x and y directions and in turn the directionality of resultant 𝑭. The external potential plugin in xml file and the ‘CellMotilitySteppable’ in python file of the CC3D code implement this active motility in cancer cells.

### Simulation details

The simulation domain is a 250 x 250 x 1 square lattice with a non-periodic boundary condition consisting of six different entities that are explained below. In the simulations, the lengths and volume dimensions are measured in terms of number of pixels or voxels, and the Monte Carlo step (MCS) is the unit of time.

*Medium (M):* This cell types constitutes of the unassigned voxels in the background on which the computational model is constructed in the simulation domain.

*Cancer cells (C1 and C2):* The Tumor consists of two types of cancer cells: Un-transitioned cancer cells (C1) which are indicated by red color and transitioned cancer cells (C2) which is indicated by yellow color. The initial single cancer cell at the center of the simulation domain has a volume of (4×4×1) voxels. Only the cancer cells in the simulation have the property to respond to chemical gradients of growth factor, grow, proliferate, and degrade ECM.

*Collagen I (CoI):* These cells mimic the fibrillar extracellular matrix (ECM) surrounding the tumor. Cancer cells degrade Collagen I and invade through it.

*C_lysed:* This cell type is depicted in an intermediate step during ECM degradation and regeneration. The dynamics of reaction-diffusion of the chemicals secreted by cancer cells allow for the degradation of CoI ECM fibers. Once the ratio of concentrations of [MMP] to [TIMP] is greater than or equal to 1.5 at the center of mass of a given *CoI* ECM fiber, it marks the onset of degradation of that fiber and it shall be converted into *C_lysed*, although retaining the shape and size of the fiber. These cells track the MCS from their individual degradation event and transform it into newly synthesized ECM cells (NC1) after 20 MCS.

*NC1:* This cell type is intended to mimic the ‘cancer-secreted matrix’, the newly synthesized ECM cells and are denoted as *NC1*. These cells are almost identical to Collagen I in their behavior and can undergo further degradation to become *C_lysed* and subsequently after 20 MCS, they will become NC1 again.

Reaction-Diffusion, growth, and proliferation equations:

The cancer cells secrete matrix metalloproteinases [MMP] and tissue inhibitors of matrix metalloproteinases [TIMP] that are involved in ECM degradation and growth factor [GF] is released from them. The chemical concentration dynamics of [MMP], [TIMP], [GF] across the simulation domain are modelled by the following reaction diffusion equations [56,57].

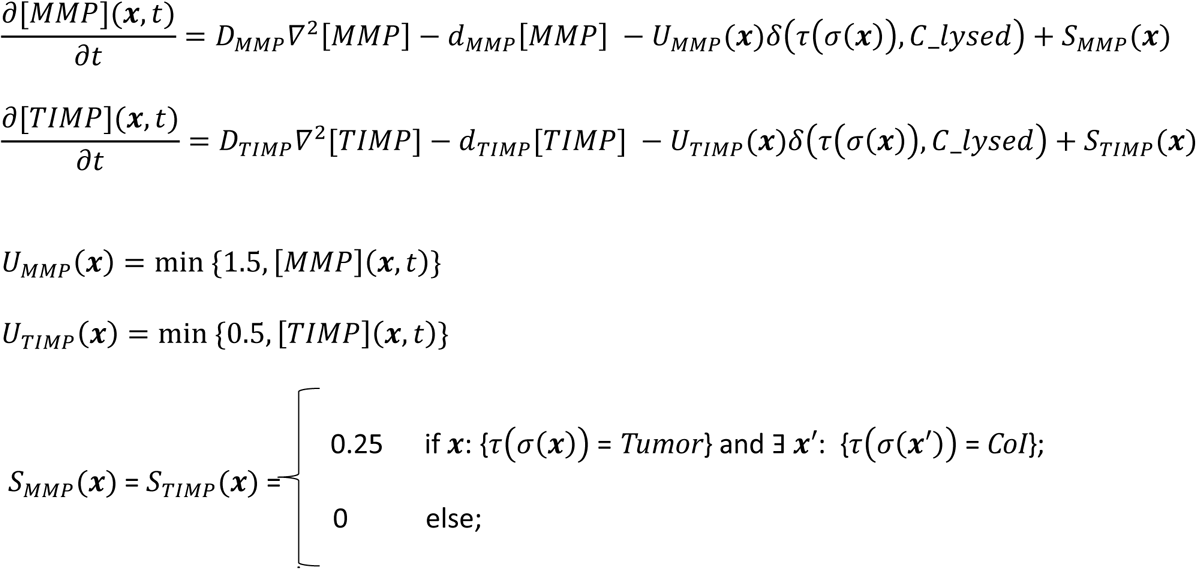

where 𝒙′ is an immediate/ first-order neighbor voxel to 𝒙 sharing a common edge, *Tumor* implies either of cancer cell types C1 or C2.

The first term in the above equations represents the diffusion term, the second being the decay term. The third term corresponds to the uptake/consumption of the chemical only by the cell type *C_lysed*. The last one being the secretion term that indicates secretion happening at only those boundary points of cancer cells that are in contact with the ECM Collagen I fibers.

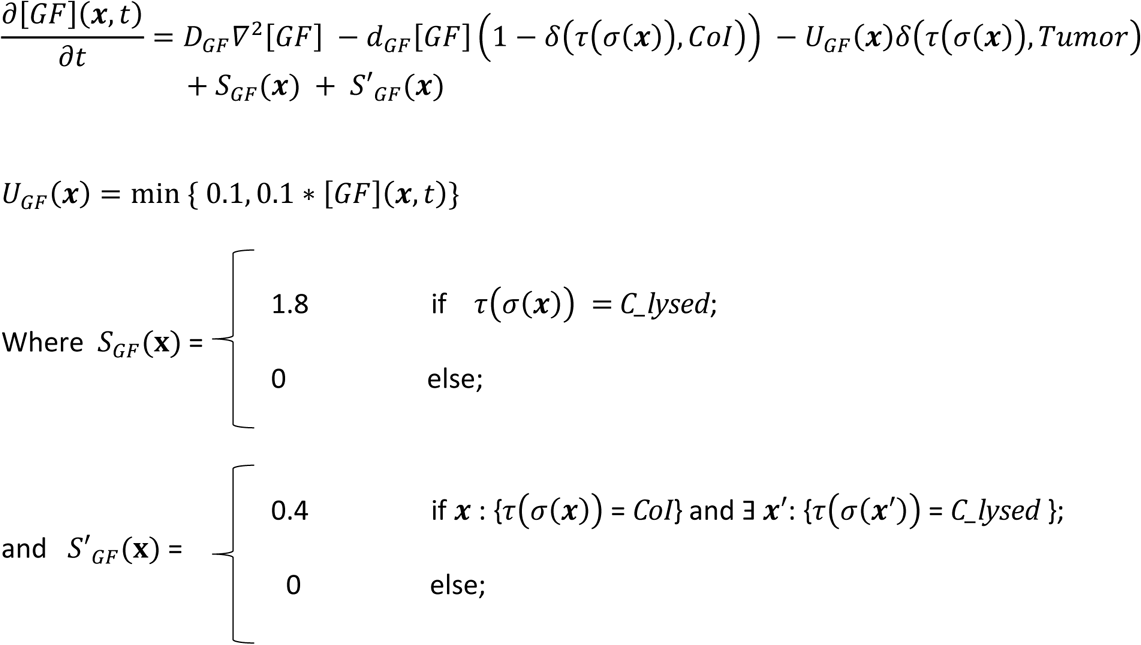

where 𝒙**’** is an immediate/ first-order neighbor voxel to 𝒙 sharing a common edge, *Tumor* implies either of cancer cell types C1 or C2.

Growth ODE for both types of cancer cells:

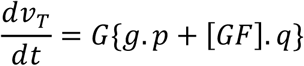

Where 𝑣_𝑇_ = Target volume of cancer cell

g = free perimeter of cancer cells available for nutrient uptake

[GF] = concentration of growth factor (GF) at center of mass of a cancer cell

p,q = constants; p = (1/30), q = (1/20)

G = Growth rate coefficient = 0.75

Once a cancer cell reaches a volume of 30 voxels, the ‘MitosisSteppable’ in the python steppables file implements cell division. The values for all the parameters described here are given in the supplementary information document and in the CC3D project code available in the GitHub link provided below.

## Statistical Analysis

All Compucell3D simulation endpoint screenshots at 8000 MCS were taken to perform image quantification analysis in MATLAB. Considering that the cellular dynamics incorporated in our model are stochastic in nature and to ensure a large sample size for statistical analysis, 60 replicates of the cases were carried out. For statistical analysis, one-way ANOVA was performed followed by the Tukey’s post hoc multiple comparisons test in GraphPad to draw inferences and conclusions. Significance (p value) is represented as *, where * ≤ 0.05, ** ≤ 0.01, *** ≤ 0.001, and **** < 0.0001.

## Data and code availability

The data and codes of the manuscript can be accessed at https://github.com/cvsp-res/EMT_CC3D

## Authors’ contribution

RB and MKJ framed the problem statement. CVSP carried out the simulations. RB, MKJ, and CVSP analyzed the results of the simulations. RB, MKJ, and CVSP wrote and edited the manuscript.

## References

[1] K. J. Cheung et al., Polyclonal breast cancer metastases arise from collective dissemination of keratin 14-expressing tumor cell clusters, Proceedings of the National Academy of Sciences 113, (2016).

[2] K. J. Cheung and S. Horne-Badovinac, Collective migration modes in development, tissue repair and cancer, Nat Rev Mol Cell Biol (2025).

[3] N. Aceto et al., Circulating Tumor Cell Clusters Are Oligoclonal Precursors of Breast Cancer Metastasis, Cell 158, 1110 (2014).

[4] A. L. Wilson, L. R. Moffitt, B. R. Doran, B. Basri, J. Do, T. W. Jobling, M. Plebanski, A. N. Stephens, and M. Bilandzic, Leader cells promote immunosuppression to drive ovarian cancer progression in vivo, Cell Rep 43, 114979 (2024).

[5] T. O. Khatib et al., TGF-β1-mediated intercellular signaling fuels cooperative cellular invasion, Cell Rep 44, 115315 (2025).

[6] J. Zhang, K. F. Goliwas, W. Wang, P. V. Taufalele, F. Bordeleau, and C. A. Reinhart-King, Energetic regulation of coordinated leader–follower dynamics during collective invasion of breast cancer cells, Proceedings of the National Academy of Sciences 116, 7867 (2019).

[7] F. Gao, G. Zhang, Y. Liu, Y. He, Y. Sheng, X. Sun, Y. Du, and C. Yang, Activation of CD44 signaling in leader cells induced by tumor-associated macrophages drives collective detachment in luminal breast carcinomas, Cell Death Dis 13, 540 (2022).

[8] S. A. Vilchez Mercedes, F. Bocci, H. Levine, J. N. Onuchic, M. K. Jolly, and P. K. Wong, Decoding leader cells in collective cancer invasion, Nat Rev Cancer 21, 592 (2021).

[9] D. Pally, D. Pramanik, S. Hussain, S. Verma, A. Srinivas, R. V. Kumar, A. Everest-Dass, and R. Bhat, Heterogeneity in 2,6-Linked Sialic Acids Potentiates Invasion of Breast Cancer Epithelia, ACS Cent Sci 7, 110 (2021).

[10] N. R. Campbell et al., Cooperation between melanoma cell states promotes metastasis through heterotypic cluster formation, Dev Cell 56, 2808 (2021).

[11] J. Konen et al., Image-guided genomics of phenotypically heterogeneous populations reveals vascular signalling during symbiotic collective cancer invasion, Nat Commun 8, 15078 (2017).

[12] T. Dutt, J. Langthasa, M. Umesh, S. Mishra, S. Bothra, K. Vidhipriya, A. Vadaparty, P. Sen, and R. Bhat, Rheological transition driven by matrix makes cancer spheroids resilient under confinement, Life Sci Alliance 8, e202402601 (2025).

[13] J. Langthasa, P. Sarkar, S. Narayanan, R. Bhagat, A. Vadaparty, and R. Bhat, Extracellular matrix mediates moruloid-blastuloid morphodynamics in malignant ovarian spheroids, Life Sci Alliance 4, e202000942 (2021).

[14] J. Yang et al., Guidelines and definitions for research on epithelial–mesenchymal transition, Nat Rev Mol Cell Biol 21, 341 (2020).

[15] E. M. Grasset et al., Triple-negative breast cancer metastasis involves complex epithelial-mesenchymal transition dynamics and requires vimentin, Sci Transl Med 14, (2022).

[16] F. Lüönd et al., Distinct contributions of partial and full EMT to breast cancer malignancy, Dev Cell 56, 3203 (2021).

[17] M. S. Brown et al., Phenotypic heterogeneity driven by plasticity of the intermediate EMT state governs disease progression and metastasis in breast cancer, Sci Adv 8, (2022).

[18] K. Hari, V. Ullanat, A. Balasubramanian, A. Gopalan, and M. K. Jolly, Landscape of epithelial– mesenchymal plasticity as an emergent property of coordinated teams in regulatory networks, Elife 11, (2022).

[19] S. Pasani, S. Sahoo, and M. K. Jolly, Hybrid E/M Phenotype(s) and Stemness: A Mechanistic Connection Embedded in Network Topology, J Clin Med 10, 60 (2020).

[20] A. Grosse-Wilde, A. Fouquier d’Hérouël, E. McIntosh, G. Ertaylan, A. Skupin, R. E. Kuestner, A. del Sol, K.-A. Walters, and S. Huang, Stemness of the hybrid Epithelial/Mesenchymal State in Breast Cancer and Its Association with Poor Survival, PLoS One 10, e0126522 (2015).

[21] S. Sahoo, S. P. Nayak, K. Hari, P. Purkait, S. Mandal, A. Kishore, H. Levine, and M. K. Jolly, Immunosuppressive Traits of the Hybrid Epithelial/Mesenchymal Phenotype, Front Immunol 12, (2021).

[22] A. Dongre et al., Direct and Indirect Regulators of Epithelial–Mesenchymal Transition–Mediated Immunosuppression in Breast Carcinomas, Cancer Discov 11, 1286 (2021).

[23] H. Agraval, K. Kandhari, and U. C. S. Yadav, MMPs as potential molecular targets in epithelial-to-mesenchymal transition driven COPD progression, Life Sci 352, 122874 (2024).

[24] D. H. Peng et al., ZEB1 induces LOXL2-mediated collagen stabilization and deposition in the extracellular matrix to drive lung cancer invasion and metastasis, Oncogene 36, 1925 (2017).

[25] A. Das, M. Monteiro, A. Barai, S. Kumar, and S. Sen, MMP proteolytic activity regulates cancer invasiveness by modulating integrins, Sci Rep 7, 14219 (2017).

[26] Y. Deng, P. Chakraborty, M. K. Jolly, and H. Levine, A Theoretical Approach to Coupling the Epithelial-Mesenchymal Transition (EMT) to Extracellular Matrix (ECM) Stiffness via LOXL2, Cancers (Basel) 13, 1609 (2021).

[27] S. Kumar, A. Das, and S. Sen, Extracellular matrix density promotes EMT by weakening cell–cell adhesions, Mol. BioSyst. 10, 838 (2014).

[28] T. Kwon, O.-S. Kwon, H.-J. Cha, and B. J. Sung, Stochastic and Heterogeneous Cancer Cell Migration: Experiment and Theory, Sci Rep 9, 16297 (2019).

[29] R. J. Murphy, P. R. Buenzli, T. A. Tambyah, E. W. Thompson, H. J. Hugo, R. E. Baker, and M. J. Simpson, The role of mechanical interactions in EMT, Phys Biol 18, 046001 (2021).

[30] F. Bocci, L. Gearhart-Serna, M. Boareto, M. Ribeiro, E. Ben-Jacob, G. R. Devi, H. Levine, J. N. Onuchic, and M. K. Jolly, Toward understanding cancer stem cell heterogeneity in the tumor microenvironment, Proceedings of the National Academy of Sciences 116, 148 (2019).

[31] S. Oliver, M. Williams, M. K. Jolly, D. Gonzalez, and G. Powathil, Exploring the role of EMT in ovarian cancer progression using a multiscale mathematical model, NPJ Syst Biol Appl 11, 36 (2025).

[32] S. Plunder, C. Danesin, B. Glise, M. A. Ferreira, S. Merino-Aceituno, and E. Theveneau, Modelling variability and heterogeneity of EMT scenarios highlights nuclear positioning and protrusions as main drivers of extrusion, Nat Commun 15, 7365 (2024).

[33] D. Pramanik, M. K. Jolly, and R. Bhat, Matrix adhesion and remodeling diversifies modes of cancer invasion across spatial scales, J Theor Biol 524, 110733 (2021).

[34] J. Xu, S. Lamouille, and R. Derynck, TGF-β-induced epithelial to mesenchymal transition, Cell Res 19, 156 (2009).

[35] C. J. David, Y.-H. Huang, M. Chen, J. Su, Y. Zou, N. Bardeesy, C. A. Iacobuzio-Donahue, and J. Massagué, TGF-β Tumor Suppression through a Lethal EMT, Cell 164, 1015 (2016).

[36] R. Y. Hapke and S. M. Haake, Hypoxia-induced epithelial to mesenchymal transition in cancer, Cancer Lett 487, 10 (2020).

[37] L. Yang et al., Pin1/ YAP pathway mediates matrix stiffness-induced epithelial– mesenchymal transition driving cervical cancer metastasis via a non-Hippo mechanism, Bioeng Transl Med 8, (2023).

[38] B. F. Matte, A. Kumar, J. K. Placone, V. G. Zanella, M. D. Martins, A. J. Engler, and M. L. Lamers, Matrix stiffness mechanically conditions EMT and migratory behavior of oral squamous cell carcinoma, J Cell Sci (2018).

[39] C.-H. Weng et al., Epithelial-mesenchymal transition (EMT) beyond EGFR mutations per se is a common mechanism for acquired resistance to EGFR TKI, Oncogene 38, 455 (2019).

[40] Y. Fujita, G. Krause, M. Scheffner, D. Zechner, H. E. M. Leddy, J. Behrens, T. Sommer, and W. Birchmeier, Hakai, a c-Cbl-like protein, ubiquitinates and induces endocytosis of the E-cadherin complex, Nat Cell Biol 4, 222 (2002).

[41] S. Sharma, H. Rani, Y. Mahesh, M. K. Jolly, J. Dixit, and V. Mahadevan, Loss of p53 epigenetically modulates epithelial to mesenchymal transition in colorectal cancer, Transl Oncol 43, 101848 (2024).

[42] Ma. C. P. Dela Cruz and P. M. B. Medina, Epithelial–mesenchymal transition (EMT) and its role in acquired epidermal growth factor receptor-tyrosine kinase inhibitor (EGFR-TKI) chemoresistance in non-small cell lung cancer (NSCLC), Cancer Pathogenesis and Therapy 3, 215 (2025).

[43] I. Pastushenko et al., Fat1 deletion promotes hybrid EMT state, tumour stemness and metastasis, Nature 589, 448 (2021).

[44] P. Dong, M. Karaayvaz, N. Jia, M. Kaneuchi, J. Hamada, H. Watari, S. Sudo, J. Ju, and N. Sakuragi, Mutant p53 gain-of-function induces epithelial–mesenchymal transition through modulation of the miR-130b–ZEB1 axis, Oncogene 32, 3286 (2013).

[45] D. Pally, S. Goutham, and R. Bhat, Extracellular matrix as a driver for intratumoral heterogeneity, Phys Biol 19, 043001 (2022).

[46] D. Pally and A. Naba, Extracellular matrix dynamics: A key regulator of cell migration across length-scales and systems, Curr Opin Cell Biol 86, 102309 (2024).

[47] D. Harjanto and M. H. Zaman, Modeling Extracellular Matrix Reorganization in 3D Environments, PLoS One 8, e52509 (2013).

[48] R. M. Crossley et al., Modeling the extracellular matrix in cell migration and morphogenesis: a guide for the curious biologist, Front Cell Dev Biol 12, (2024).

[49] O. Ilina et al., Cell–cell adhesion and 3D matrix confinement determine jamming transitions in breast cancer invasion, Nat Cell Biol 22, 1103 (2020).

[50] M. M. Martino and J. A. Hubbell, The 12th–14th type III repeats of fibronectin function as a highly promiscuous growth factor-binding domain, The FASEB Journal 24, 4711 (2010).

[51] J. M. Wells, A. Gaggar, and J. E. Blalock, MMP generated matrikines, Matrix Biology 44–46, 122 (2015).

[52] J. D. Mott and Z. Werb, Regulation of matrix biology by matrix metalloproteinases, Curr Opin Cell Biol 16, 558 (2004).

[53] J. Winkler, A. Abisoye-Ogunniyan, K. J. Metcalf, and Z. Werb, Concepts of extracellular matrix remodelling in tumour progression and metastasis, Nat Commun 11, 5120 (2020).

[54] F. Graner and J. A. Glazier, Simulation of biological cell sorting using a two-dimensional extended Potts model, Phys Rev Lett 69, 2013 (1992).

[55] M. H. Swat, G. L. Thomas, J. M. Belmonte, A. Shirinifard, D. Hmeljak, and J. A. Glazier, Multi-Scale Modeling of Tissues Using CompuCell3D, in (2012), pp. 325–366.

[56] M. Scianna and L. Preziosi, Multiscale Developments of the Cellular Potts Model, Multiscale Modeling & Simulation 10, 342 (2012).

[57] C. Giverso, M. Scianna, L. Preziosi, N. Lo Buono, and A. Funaro, Individual Cell-Based Model for In-Vitro Mesothelial Invasion of Ovarian Cancer, Math Model Nat Phenom 5, 203 (2010).

