## Supplementary information for "Multiscale modeling predicts dependence of mesenchymally transitioned tumor niche fitness on cell-cell and cell-matrix adhesions"

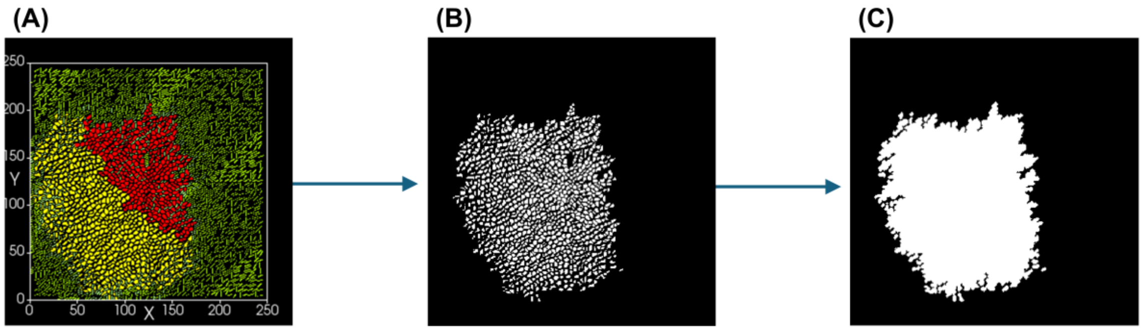


**Figure S1: Image analysis and quantification of end point of simulation**

(A) Tumor at end point of simulations (8000 MCS) with un-transitioned niche (red) and transitioned niche (yellow) with complete EMT.

(B) Grayscale image consisting of only cancer cells of the end point of simulation run.

(C) Grayscale image after application of image morphological operations (strel, imdilate, imfill, bwareaopen in MATLAB) depicting the area of largest cell cluster (ALC) and the remaining small cell clusters at the periphery contributing to number of disconnected cell clusters (NDC).

#### Adhesion strengths for cancer cells corresponding to Fig. 1 & 2 (Complete EMT):

In CC3D, adhesion strengths are specified in terms of contact energies. Contact energies and adhesion strengths are inversely related, implying low contact energies correspond to high adhesion strengths and vice versa. The contact energies of un-transitioned (red) cells correspond to the control 'No Transition' cases. Collagen I here is same as extracellular matrix (ECM) referred in the main manuscript.

Intra-niche contact energy among the un-transitioned (red) cells: 46

Intra-niche contact energy among the transitioned (yellow) cells: **92**

Inter-niche contact energy between a red and a yellow cell: **92**

Cell-Collagen I contact energy for un-transitioned cells: 46

Cell-Collagen I contact energy for transitioned cells: **23**

**Adhesion strengths for cancer cells corresponding to Fig. 3 (High cell-ECM adhesion transition):**

Intra-niche contact energy among the un-transitioned (red) cells: 46

Intra-niche contact energy among the transitioned (yellow) cells: 46

Inter-niche contact energy between a red and a yellow cell: 46

Cell-Collagen I contact energy for un-transitioned cells: 46

Cell-Collagen I contact energy for transitioned cells: **23**

**Adhesion strengths for cancer cells corresponding to Fig. 4 (Low inter- and intra-niche adhesion transition):**

Intra-niche contact energy among the un-transitioned (red) cells: 46

Intra-niche contact energy among the transitioned (yellow) cells: **92**

Inter-niche contact energy between a red and a yellow cell: **92**

Cell-Collagen I contact energy for un-transitioned cells: 46

Cell-Collagen I contact energy for transitioned cells: 46

**Adhesion strengths for cancer cells corresponding to Fig. 5 (High cell-ECM adhesion and low inter- niche adhesion transition):**

Intra-niche contact energy among the un-transitioned (red) cells: 46

Intra-niche contact energy among the transitioned (yellow) cells: 46

Inter-niche contact energy between a red and a yellow cell: **92**

Cell-Collagen I contact energy for un-transitioned cells: 46

Cell-Collagen I contact energy for transitioned cells: **23**

**Adhesion strengths for cancer cells corresponding to Fig. 6 (High cell-ECM adhesion and low intra- niche adhesion transition):**

Intra-niche contact energy among the un-transitioned (red) cells: 46

Intra-niche contact energy among the transitioned (yellow) cells: **92**

Inter-niche contact energy between a red and a yellow cell: 46

Cell-Collagen I contact energy for un-transitioned cells: 46

Cell-Collagen I contact energy for transitioned cells: **23**

**Other key parameters of the model that are fixed in all simulations:**

| Parameter | Symbol/notation | value |
|---|---|---|
| Initial target volume of cancer cells | $v_{T0}$ | 24 voxels |
| Lambda volume of cancer cells | $\lambda_v$ | 20 |
| Target surface of cancer cells | $s_T$ | 20 voxels |
| Lambda surface of cancer cells | $\lambda_s$ | 2 |
| Target volume of all ECM fibers | $(v_T)_{\text{ECM}}$ | 12 voxels |
| Lambda volume of all ECM fibers | $(\lambda_v)_{\text{ECM}}$ | 20 |
| Chemotaxis strength | $\lambda_{chem}$ | 100 |
| Diffusion constant of [MMP] | $D_{MMP}$ | 0.01 |
| Decay constant of [MMP] | $d_{MMP}$ | 0.003 |
| Diffusion constant of [TIMP] | $D_{TIMP}$ | 0.035 |
| Decay constant of [MMP] | $d_{TIMP}$ | 0.003 |
| Diffusion constant of [GF] | $D_{GF}$ | 0.05 |
| Decay constant of [GF] | $d_{GF}$ | 0.001 |
| MMP-TIMP RD cooperativity threshold | $\{[MMP]/[TIMP]\}_{\text{threshold}}$ | 1.5 |
| Coeff. of cancer cell free perimeter | p | 1/30 |
| Coeff. of [GF] concentration term | q | 1/20 |
| Growth rate coefficient | G | 0.75 |

### Supplementary videos

**Video S1:** Simulation run of transitioned niche having higher cell-ECM adhesion **confined** by the un-transitioned niche.

**Video S2:** Simulation run of transitioned niche having higher cell-ECM adhesion **unconfined** by the un-transitioned niche.
